# Microglial synaptic pruning in the nucleus accumbens during adolescence sex-specifically influences splenic immune outcomes

**DOI:** 10.1101/2023.05.03.539317

**Authors:** J. M. Kirkland, Ishan Patel, Monali S. Ardeshna, Ashley M. Kopec

**Affiliations:** Department of Neuroscience and Experimental Therapeutics, Albany Medical College

## Abstract

Strong social support promotes a variety of positive health outcomes in humans and rodent models, while social isolation in rodents shortens lifespan, perceived social isolation (i.e. loneliness) can increase mortality by up to 50% in humans. How social relationships lead to these drastic health effects is unclear, but may involve modulation of the peripheral immune system. The reward circuitry of the brain and social behaviors undergo a critical period of development during adolescence. We published that microglia-mediated synaptic pruning occurs in the nucleus accumbens (NAc) reward region during adolescence to mediate social development in male and female rats. We hypothesized that if reward circuitry activity and social relationships directly impact the peripheral immune system, then natural developmental changes in the reward circuitry and social behaviors during adolescence should also directly impact the peripheral immune system. To test this, we inhibited microglial pruning in the NAc during adolescence, and then collected spleen tissue for mass spectrometry proteomic analysis and ELISA validation. We found that the global proteomic consequences of inhibiting microglial pruning in the NAc were similar between the sexes, but target-specific examination suggests that NAc pruning impacts Th1 cell-related immune markers in the spleen in males, but not females, and broad neurochemical systems in the spleen in females, but not males.

Please note, if this preprint will be pushed further to publication it will not be by me (AMK), as I am leaving academia. So, I’m going to write more conversationally.

## INTRODUCTION

Social behaviors are integral to health and wellness across the lifespan. Strong social support promotes a variety of positive health outcomes in humans and rodent models, while social isolation in rodents shortens lifespan, perceived social isolation (i.e. loneliness) can increase mortality by up to 50% in humans [1, 2]. How social relationships lead to these drastic health effects is unclear, but may involve modulation of the peripheral immune system [3, 4]. The impact of peripheral immune signaling on neural and behavioral outcomes is now well-established, but social context also regulates peripheral immune responses. For example, in macaques, high vs. low social status reliably predicts expression of immune gene networks in blood, and these networks dynamically change if social status is manipulated [5]. Social behaviors are supported in part by the ‘reward’ circuitry of the brain [4, 6, 7]. Asya Rolls and colleagues have elegantly shown that artificially activating the reward circuitry can *on its own* enhance pathogen and tumor-induced immune responses in the peripheral immune system, including in the largest immune organ of the body, the spleen [8, 9]. These data suggest that the reward circuitry neural activity engaged during social behaviors can concurrently impact the body’s response to immune challenges.

The reward circuitry of the brain and social behaviors undergo a critical period of development during adolescence. One neurodevelopmental mechanism that appears to be common across brain regions and developmental periods is the requirement for synaptic pruning to produce mature neural communication and circuits. In many cases, this is mediated via immune signaling: microglia, the resident immune cells of the brain, express complement receptor 3 (CR3/CD11b) which binds to its ligand C3, a phagocytic ‘tag’ that associates with weak or inactive synapses designated for pruning [10-12]. We published that microglia-C3-mediated synaptic pruning also occurs in the nucleus accumbens (NAc) reward region during adolescence to mediate social development in male and female rats [12]. In males, NAc pruning occurred between early and mid adolescence (postnatal day (P)30-40) to mediate social development, while in females NAc pruning occurred between pre- and early adolescence (P20-30) to mediate social development [12]. Adolescence is also a period during which immune responses dynamically change in macaques, humans, and rats [13-16]. In humans, males mount larger immune responses than females prior to adolescence, which reverses after adolescence [13]. And, similar to our observations in the NAc, immune responses in the spleen are reported to mature during pre-adolescence in female rats (P21) and mid-adolescence in males (P42) [17]. We hypothesized that if reward circuitry activity and social relationships directly impact the peripheral immune system in adulthood, then natural developmental changes in the reward circuitry and social behaviors during adolescence should also directly impact the peripheral immune system. To test this, we inhibited microglial pruning in the NAc during each sex’s pruning period, and then collected spleen tissue for mass spectrometry proteomic analysis and ELISA validation. We found that the global proteomic consequences of inhibiting microglial pruning in the NAc were similar between the sexes, but target-specific examination suggests that NAc pruning impacts immune markers in the spleen in males, but not females, and broad neurochemical systems in the spleen in females, but not males.

## METHODS

### Animal Care

Adult male and female Sprague-Dawley rats were purchased to be breeding pairs (Harlan/Envigo). Litters were culled to a maximum of 12 pups between postnatal day (P)2-5, and pups were weaned into same-sex housing in pairs or triplets at P21. At least 3 litters were represented in each experimental group. Rats were housed in conventional cages on cellulose bedding with ad libitum access to food and water. Colonies were maintained on a 12:12 light:dark cycle (lights on a 07:00) in a temperature- (20-24°C) and humidity- (35-55 RH) controlled room. Cages were changed twice weekly. All experiments were approved by the Institutional Animal Care and Use Committee at Albany Medical College.

### Animal Model

Rats underwent surgical intervention at either P22 (females) or P30 (males) according to our previously published sex-specific NAc pruning periods [12]. Rats were anesthetized under isoflurane (2-3%) and fastened into the stereotactic apparatus. The scalp was cut midsagitally and holes drilled to target NAc bilaterally. NIF (5µg/µL) or Vehicle (sterile PBS) was infused into the NAc using a Hamilton syringe at 10° using sex-specific coordinate and volume parameters: AP +2.25mm, ML ±2.5mm, DV −5.75mm, 300nL for P30 males and AP +2.7mm, ML ±2.4 mm, DV -5.55mm, 250nL for P22 females. One depth was reached, the syringe remained in place for 1 min before infusion (50nL/min), and for 5 mins post-infusion. The wound was closed with surgical staples and treated topically with anti-bacterial ointment and Bupivicaine. Ketofen (5mg/kg) was administered subcutaneously after surgery, and once daily for two days after surgery.

### Sample Collection and Preparation

Rats were euthanized 8 days after surgery, a time point we have published that molecular and behavioral effects of NIF treatment are evident in both sexes [12], and perfused transcardially with saline. Spleens were extracted, rapidly frozen, and homogenized in RIPA buffer with protease and phosphatase inhibitors.

Homogenate from two sex- and manipulation-matched rats were pooled into one sample for proteomic analysis carried out by Creative Proteomics (https://www.creative-proteomics.com/).

### Mass Spectrometry (performed by Creative Proteomics)

200µg protein per sample was precipitated with cold acetone. Pellets were dissolved in 2M urea and denatured with 10mM DTT for 1hr at 56°C followed by alkylation with 50nM IAA for 1hr at room temperature in the dark.

Ammonium bicarbonate was added to a final concentration of 50mM, pH 7.8, and trypsin was added to the sample for digestion for 15 hrs at 37°C. Peptides were purified with a C18 SPE column and lyophilized to near dryness. Peptides were resuspended in 20µL of 0.1% formic acid before LC-MS/MS analysis. 1µg of sample was loaded onto a nanoLC-MS/MS platform. The full scan was performed between 300-1650 m/z at a resolution of 60,000 at 200 m/z. The gain was set to 3e6. The MS/MS scan operated in Top 20 mode with 15,000 at 200 m/z resolution, 1e5 automatic gain control target, 19ms maximum injection time, 28% normalized collision energy, 1.4 Th isolation window, unassigned, 1, >6 charge state exclusion, and 30 sec dynamic exclusion. Raw MS files were aligned with the rat protein database using Maxquant (1.6.1.14). Protein modifications were carbamidomethylation (fixed), oxidation (variable), the enzyme specificity was set to trypsin, the maximum missed cleavage was set to 2, the precursor ion mass tolerance was set to 10 ppm, and the MS/MS tolerance was 0.5 Da.

### Data Analysis and Statistics

Data from Creative Proteomics were quality controlled in-house: any proteins that had more than one sample in which they were not detected (0 intensity) within the group of 3-4 total samples was excluded from analysis. In proteins with just one non-detected sampled, the average of the other 2-3 samples was used to replace the 0. *t*-tests were performed between NIF and Vehicle groups for each sex. In the case of duplicate protein IDs, the ID corresponding to the smallest p-value was retained and the others were excluded from analysis. Protein groups that were significantly changed by NIF treatment (*p*<0.05, no correction for multiple comparisons) were assessed for global proteomic summary, including Pearson’s correlation of NIF-induced changes in all proteins between the sexes (*Fig. 1*). Benjamini-Hochberg analyses (FDR = 0.1) were performed to determine the proteins significantly changed by NIF treatment after correction for multiple comparisons (*Fig. 2*). Protein concentrations of remaining unpooled homogenate was determined with BCA assays following the manufacturer’s protocols. ELISAs were conducted per the manufacturer’s protocols by loading 2000µg total protein per sample. ACh and TH ELISAs were purchased from Novus Biologicals (#NBP2-66389 and #NBP3-06922, respectively). NA ELISA was purchased from LS Bio (#LS-F28027-1). IL-2, TNFα, and IFNγ ELISAs were purchased from R&D Systems (#R2000, #RTA00, and #RIF00, respectively). ELISA data were analyzed with *t*-tests for each sex. For all analyses, statistical significance is defined as *p*<0.05. Four datapoints across the sex ELISAs were identified as outliers via Grubb’s test (α=0.05) and excluded from statistical analysis. All statistics were performed in GraphPad Prism 9.5.1

**Fig. 1:**
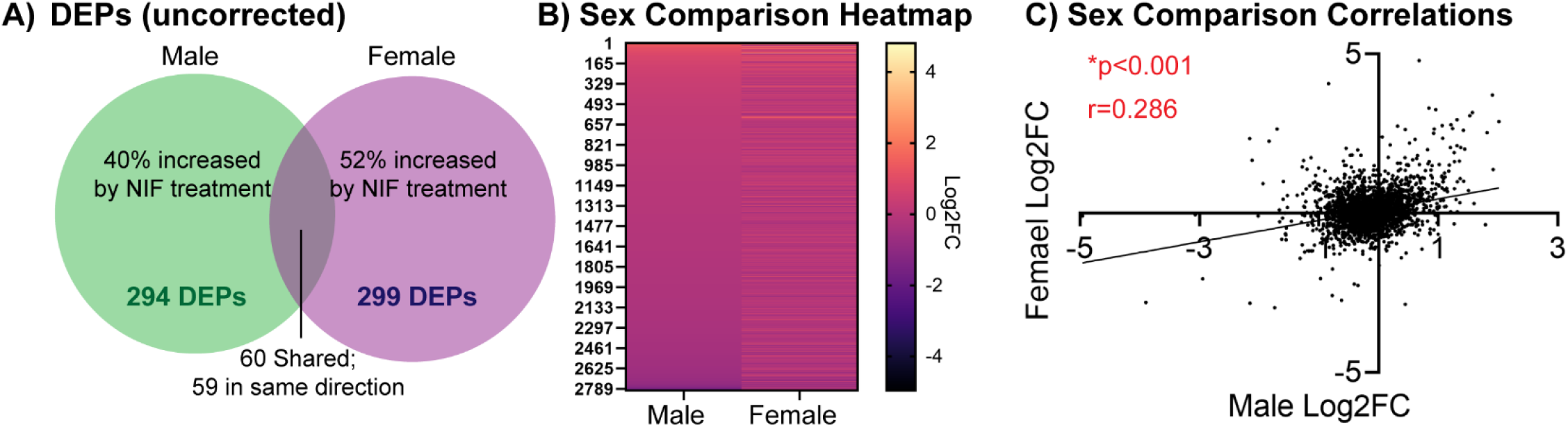
Inhibition of microglial synaptic pruning in the NAc induces similar global proteomic outcomes in male and female rat spleen. Male and female rats were stereotactically injected with NIF or Vehicle bilaterally into the NAc at each sex’s respective pruning period (P30 in males, P22 in females). Eight days later, rats were euthanized and spleens frozen and sent for proteomic analysis. Two sex- and treatment-matched rats’ NAc samples were pooled into one biological replicate to ensure enough protein yield. NAc samples were processed via label-free nanoLC-MS/MS mass spectrometry. **(A)** After quality control, 2930 and 2933 total proteins were identified in male and female spleen, respectively, of which 294 and 299, respectively, were significantly changed (differentially expressed proteins; DEPs) at α=0.05 uncorrected for multiple comparisons. Of these, 40% were upregulated by NIF-mediated inhibition of NAc pruning in males, and 52% were upregulated in females. Sixty DEPs were shared between the sexes, of which 59 were regulated in the same direction (see **Supp. Table 1**). **(B)** Of all proteins identified, 2800 were shared between the sexes. Plotting the fold change of each protein in both sexes suggests that inhibiting NAc pruning in males and females result in similar regulation of the proteomic landscape in the spleen, e.g. a protein upregulated in males is also likely to be upregulated in females. **(C)** To assess this quantitatively, we examined the correlation in splenic protein expression changes induced by NAc pruning inhibition between males and females, and found a significant positive correlation, indicating that our visual observation in *B* was the case. *n*=3-4 biological replicates/sex/condition. \**p*<0.05.

**Fig. 2:**
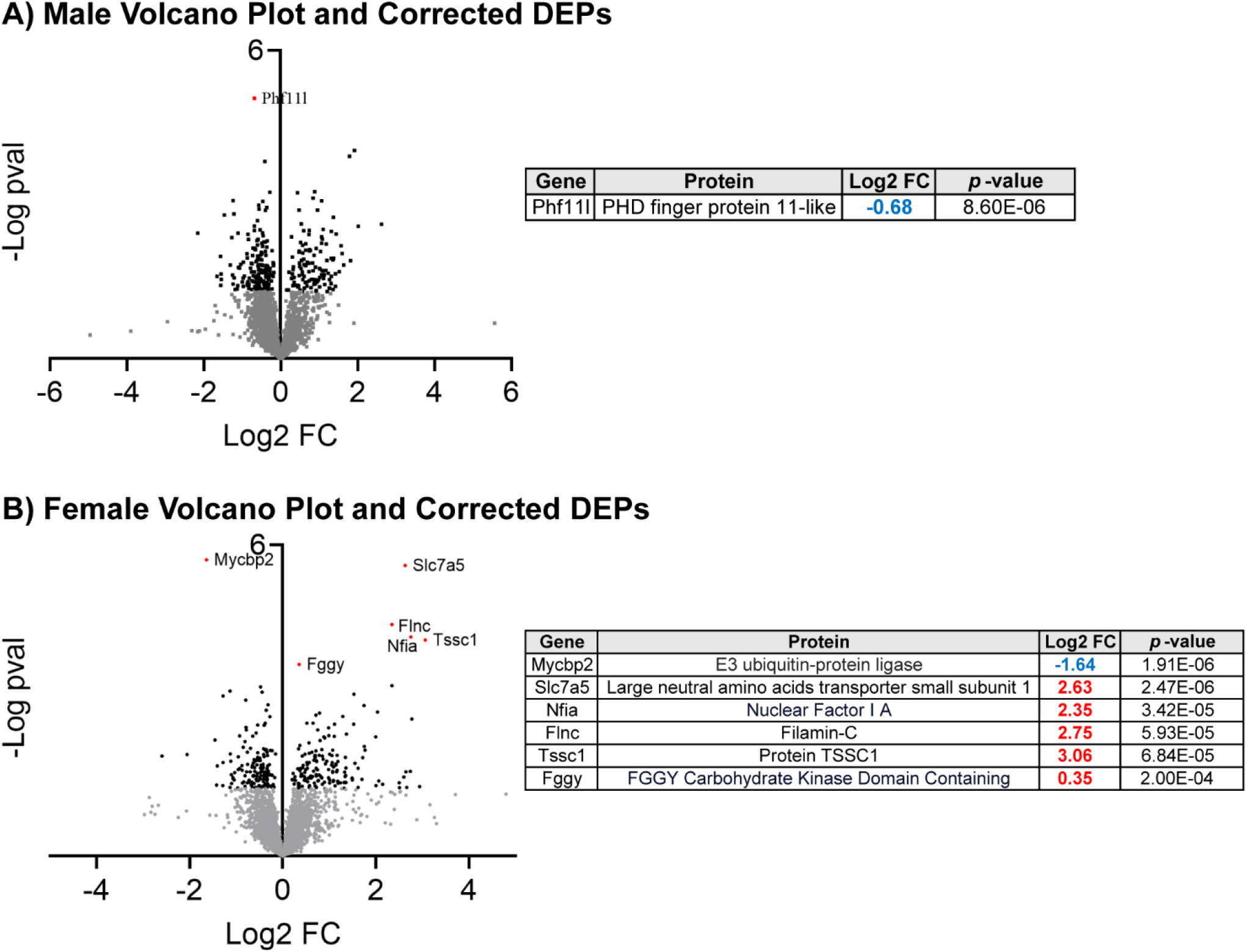
Sex-specific DEPs in the spleen induced by inhibition of microglial synaptic pruning in the NAc. Volcano plots depict each identified splenic protein in each sex, with gray dots indicating non-significant (α=0.05 uncorrected for multiple comparisons), black dots indicating significant DEPs (α=0.05 uncorrected for multiple comparisons), and red, labeled dots indicating significant DEPs after correction for multiple comparisons (Benjamini-Hochberg; FDR=0.1). Proteins in Quadrant 1 are upregulated, and proteins in Quadrant II are downregulated. **(A)** In males, only 1 DEPs passed multiple comparisons correction, which is identified in the table to the right. PHf11l was downregulated (blue FC value). **(B)** In females, 6 DEPs passed multiple comparisons correction, which are identified in the table to the right. One DEP was upregulated and the other 5 were upregulated (red FC value). *n*=3-4 biological replicates/sex/condition.

## RESULTS

Because the male and female NAc pruning period is sex-specific [12], we performed surgeries to inhibit NAc pruning at different ages (P22 in females, P30 in males). Thus, we cannot directly compare the sexes.

However, we can compare patterns induced by inhibiting pruning in the NAc between the sexes. We will begin with a global summary of these patterns before examining unique outcomes in each sex.

### Inhibition of microglial synaptic pruning in the NAc induces similar global proteomic outcomes in male and female rat spleen

In total, 2930 and 2933 total proteins were identified in the spleen via label free mass spectrometry after NIF-or Vehicle-treatment of the NAc in adolescent males and females, respectively. Of these, there were 294 differentially expressed proteins (DEPs; uncorrected for multiple comparisons) in males and 299 DEPs in females. Sixty of these proteins were differentially expressed in both sexes, and of these only one protein was *not* regulated in the same direction (**Fig. 1A; Supp. Table 1**). A heatmap comparison of the adolescent NAc NIF-induced fold change in all splenic proteins suggested that male and female outcomes may be similar (**Fig. 1B**), which was confirmed with a significant positive correlation (*r*=0.286, *p*<0.001; **Fig. 1C**).

### Sex-specific DEPs in the spleen induced by inhibition of microglial synaptic pruning in the NAc

As discussed in *Fig. 1*, NIF-mediated inhibition of microglial pruning in the NAc appeared to regulate male and female splenic protein expression in the same way. To better understand the unique consequences of NAc pruning in each sex, we conducted Benjamini-Hochberg adjustment for multiple comparisons (**Fig. 2**), which revealed 1 splenic DEP in males and 6 splenic DEPS in females that were still significantly regulated by adolescent NIF treatment in the NAc. In males, Phf11l was significantly downregulated in the spleen after inhibiting pruning in the NAc. In females, Mycbp2 was significantly downregulated and Slc7a5, Nfia, Finc, Tssc1, and Fggy were significantly upregulated after inhibiting pruning in the NAc. There was no meaningful pathway enrichment for either sex using either corrected or uncorrected DEP lists at the α = 0.05 level (*data not shown*).

### Inhibition of microglial synaptic pruning in the NAc increases IL-2 expression in the spleen in males, but not females

We could not discern a pattern or testable consequence of the female-specific splenic DEPs shown in *Fig. 2* to further assess with target-specific assays. However, the lone male spleen DEP that was significantly regulated by inhibiting pruning in the NAc, Phf11l, did provide an opportunity for further assessment. Phf11l, Plant homeodomain finger protein 11-like is a transcriptional regulator of interleukin-2 (IL-2) and interferon-γ(IFNγ) widely associated with Th1 immune cells [18, 19]. We thus used unpooled samples left from our mass spectrometry analyses to quantify these cytokines and another common immune cytokine, tumor necrosis factor-α(TNFα; **Fig. 3**). Inhibiting pruning in the NAc during adolescence increased splenic IL-2 expression, in males only (males: *t*(15)=2.417, *p*=0.029; females: *t*(13)=0.157, *p*=0.878). NIF treatment did not statistically significantly regulate either IFNγ(males: *t*(15)=1.383, *p*=0.187; females: *t*(12)=1.439, *p*=0.176) or TNFα (males: *t*(14)=1.904, *p*=0.078; females: *t*(13)=0.298, *p*=0.771) in the spleen of either sex.

**Fig. 3:**
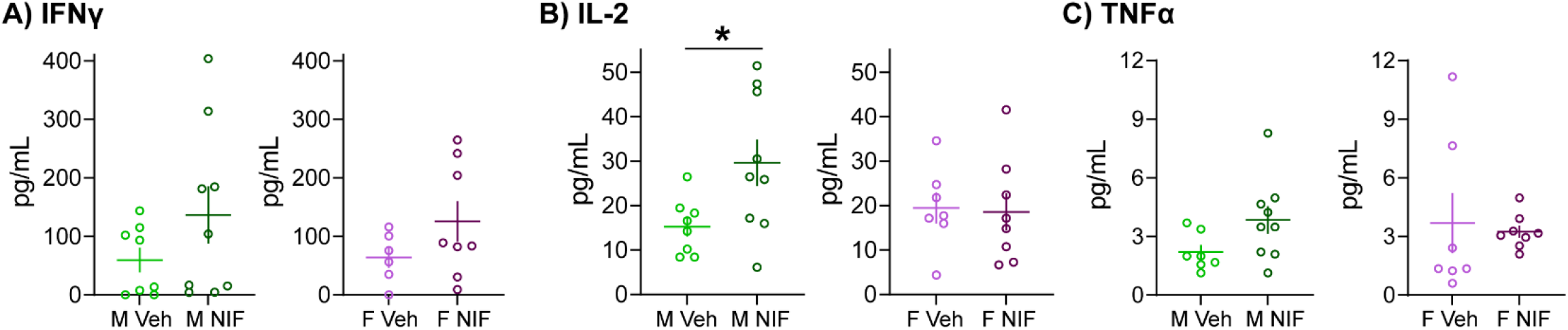
Inhibition of microglial synaptic pruning in the NAc increases IL-2 expression in the spleen in males, but not females. In remaining tissue (unpooled) that underwent proteomic analysis (*Figs. 1, 2*) we conducted ELISAs to better understand splenic changes in response to manipulating NAc development via measurement of cytokines important for immune function: interferon-γ (IFNγ), interleukin-2 (IL-2), and tumor necrosis factor α (TNFα). Inhibiting pruning in the NAc **(B)** increased splenic IL-2 levels in males, but not females. **(A)** IFNγ and **(C)** TNFα were not changed by NIF-treated in either sex. In each histogram, horizontal lines are average and vertical lines are standard error of the mean. *n*=7-8/sex/condition. \**p*<0.05.

**Fig. 3:**
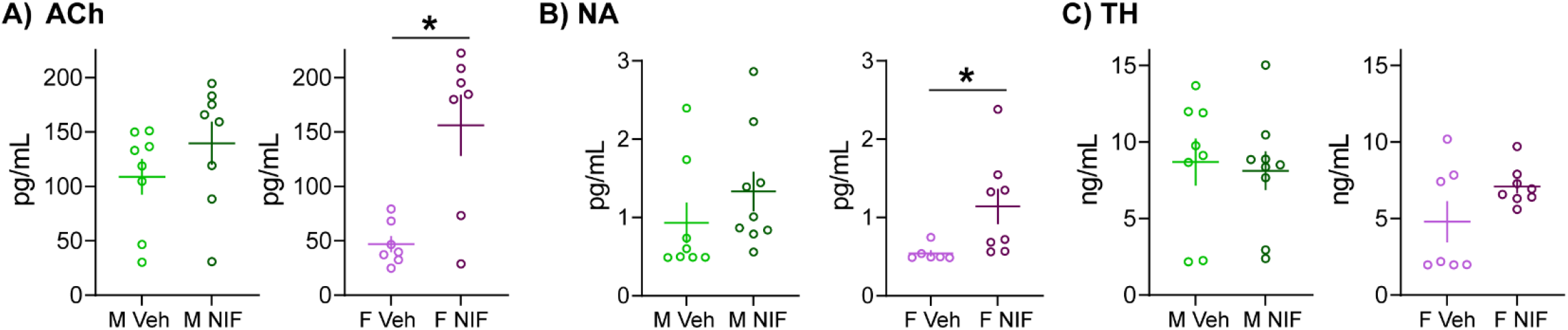
Inhibition of microglial synaptic pruning in the NAc alters neurochemical systems in the spleen in females, but not males. In remaining tissue (unpooled) that underwent proteomic analysis (*Figs. 1, 2*) we conducted ELISAs to better understand splenic changes in response to manipulating NAc development via measurement of markers of neurochemical systems broadly important for splenic function: acetylcholine (ACh), noradrenaline (NA), and tyrosine hydroxylase (TH). Inhibiting pruning in the NAc incresaed **(A)** ACh and **(B)** NA levels in the spleen in females, but not males. NIF treatment did not statistically significantly change TH levels in either sex. In each histogram, horizontal lines are average and vertical lines are standard error of the mean. *n*=7-8/sex/condition. \**p*<0.05.

### Inhibition of microglial synaptic pruning in the NAc alters neurochemical systems in the spleen in females, but not males

Finally, we tested whether neurochemical systems more broadly important for splenic immune function are altered by NAc pruning during adolescence (**Fig. 4**). Acetylcholine (ACh) signaling plays an important role in the anti-inflammatory reflex mediated indirectly via the parasympathetic vagus nerve [20], noradrenaline (NA) is the primary neurochemical system used by the sympathetic nervous system and the only direct source of innervation to the spleen [21, 22], and tyrosine hydroxylase is a rate-limiting enzyme in the synthesis of noradrenaline and dopamine that is present in innervating neural terminals and some immune cells [22]. Inhibiting pruning in the NAc during adolescence increased ACh and NA expression in the spleens of females, but not males (ACh (males: *t*(14)=1.195, *p*=0.252; females: *t*(12)=3.756, *p*=0.003); NA (males: *t*(15)=1.108, *p*=0.285; females: *t*(12)=2.268, *p*=0.043)). There was no statistically significant change in TH expression in the spleen resulting from NAc NIF treatment (males: *t*(15)=0.291, *p*=0.775; females: *t*(13)=1.715, *p*=0.110).

## DISCUSSION

In this set of studies, we determined how microglia-mediated synaptic pruning in the NAc during adolescence impacts the proteomic landscape in the spleen in male and female rats. We observed that globally, there were similar changes occurring in male and female spleen in response to inhibition of NAc pruning, despite NAc pruning occurring at different adolescent ages in each sex (P22 in females, P30 in males). However, the sets of DEPs regulated by pruning were unique to each sex, which led us to examine immune and splenic neurochemistry in more detail. Inhibiting NAc pruning during adolescence in males (but not females) increased expression of IL-2 in the spleen, while in females (but not males) inhibiting NAc pruning during adolescence increased ACh and NA expression in the spleen.

### The reward circuitry directly modulates the peripheral immune system

Since the recognition of social withdrawal and anhedonia reliably accompanies sickness [23], the field has appreciated that neural activity within the reward circuitry and associated reward-related behaviors are influenced by the status of the peripheral immune system. In humans, experimentally-induced peripheral immune activation too low to induce overt sickness can change fMRI BOLD signal in the NAc and is associated with increased preference for familiar social stimuli like family members over novel social stimuli [24].

Monocyte trafficking from the spleen to the brain is also causally implicated in depression-like behaviors in mice [25, 26]. However, the brain is not a helpless bystander. Early work with dopaminergic lesions in the brain indicated that different brain regions control different aspects of peripheral immunity. In fact, dopaminergic denervation of the NAc was reported to reduce splenic immune responses to immune challenge [27]. More recent work has now clearly shown that experimental activation of the ventral tegmental area of the reward circuitry enhances immune responses to future immune challenges [8, 9]. Initially, the prevailing theory was that the brain exerts splenic control via the parasympathetic vagus nerve. That has only turned out to be partially true. Although the vagus nerve is critical for the ‘cholinergic anti-inflammatory reflex’, which reduces pro-inflammatory signaling in the spleen, it is generally accepted that the vagus nerve does not directly innervate the spleen. Instead, the vagus nerve innervates the sympathetic celiac ganglion, which in turn relays activity to acetylcholine-expressing T cells in the spleen [28, 29]. The only direct innervation to the spleen is noradrenergic sympathetic innervation from the splenic nerve [21, 22]. Consistent with this, several studies have shown that neural control over the spleen requires the sympathetic nervous system, and not the parasympathetic nervous system [8, 25]. There are several splenic nerve collaterals, which recently have been reported to respond differentially to immune challenges [30]. I would be willing to bet that different central neural activity corresponds with each splenic nerve collateral.

Several studies have shown that prolonged social stress, including prolonged social stress during adolescence [31], results in peripheral immune dysfunction. These studies are the first to show that a *single* developmental mechanism, synaptic pruning, within a *single* brain region, the NAc, significantly impacts the spleen. However, it is still possible that the NAc is a red herring. Inhibiting microglial pruning in the NAc not only changes the local NAc environment, it changes social behavior by increasing social play acutely during adolescence [12]. Because social play is mediated by many different brain regions, it is possible that the splenic effects we observe are in fact caused by altered neural activity in other brain regions, and less directly reliant on the NAc. It is also important to note that while we manipulated microglial pruning in the NAc in both sexes, the splenic outcomes were not equivalent. Moreover, the NAc pruning period is not the same in both sexes, nor is the pruning target the same in both sexes [12, 32]. Males and females are very different. They should be studied in parallel (and no, the females aren’t more “variable” [33] so stop with the nonsense and include females in your studies). What exactly the neural to splenic link is that mediates the effects reported herein remains to be uncovered, but the takeaway is important regardless: adolescent neural/behavioral development influences the peripheral immune system.

### Functional implications of sex-specific splenic regulation during adolescence

While only one splenic DEP was significantly regulated by inhibiting microglial pruning in the NAc in males after correcting for multiple comparisons, it turned out to be lead us down an informative path. Phf11 is associated with allergy atopic dermatitis in children [34]. It is expressed at a higher level in Th1 cells than in Th2 cells [19], and its expression can be modulated by TLR3 activation [35]. In T cells, Phf11 shuttles from the cytoplasm to the nucleus and acts a transcriptional co-activator with NF_κ_B to regulate Th1 effector genes [18]. Although both IFNγ and IL-2 are Th1 effector genes, overexpression of Phf11 reportedly increased transcription of IFNγ, but not IL-2, while knock down of Phf11 reduced transcription in both genes [18]. Phf11 is also important in B cells for responses to IgE [36], but there are slightly less data (that I could find) regarding its role in B cells. Our data indicate that inhibiting NAc pruning during adolescence reduces Phf11 levels in the spleen in males, but not females. We also found that inhibiting NAc pruning increased IL-2 levels, but not (statistically significantly) IFNγ or TNFα levels in the spleen in males, but not females. This means that *natural* NAc pruning during adolescence serves to *increase* splenic Phf11 and IL-2 levels in males, but not females. This could be accomplished by Th1 cells (or other immune cells) increasing experiencing an increase in Phf11, which then increases IL-2 via its transcriptional upregulation. It could also be accomplished by just having a higher concentration of Th1 cells in the spleen. T cell egress from the spleen is reported [37, 38], so perhaps there is a cessation of brain to sympathetic nervous system neural signals that are responsible for splenic T cell egress during adolescence. If I were to continue these studies, I would first check whether NAc pruning changes the ratio of immune cell populations present in the spleen. The answer, and why it is male-specific, is a question for a future set of studies.

In females, inhbiting microglial pruning in the NAc during adolescence resulted in more significantly regulated splenic DEPs after multiple comparisons correction, but none of the DEPs (to my non-immunologist eyes) were clearly leading us to test other immune factors as was the case in the males. However, because of the opposing neurochemical systems at play in the spleen, ACh promoting anti-inflammation and NA promoting inflammation, we sought to test whether there were changes in these more broad systems. Inhibiting microglial pruning in the NAc increased ACh and NA levels in the spleen in females, but not males. That means that *natural* NAc pruning during adolescence serves to *decrease* these systems in the spleen in females, but not males. In the case of NA, it is possible that sympathetic neural innervation to the spleen is being refined during adolescence. TH is loosely considered to be a marker of neural innervation in the spleen [22], but the reality is that many immune cells also express TH. So, the fact that TH doesn’t change doesn’t necessarily rule out the possibility that increased NA is due to increased innervation. Synaptophysin or another truly unique pre-synaptic neural marker would better assess that hypothesis. It could also be the case that the existing neural innervation does not change, but is more active. Several studies indicate that neural activity in the brain is translated to splenic outcomes via activity from the sympathetic splenic nerve [8, 25]. More neural activity in brain regions that directly impact the splenic nerve would thus make sense, and it is a good bet that inhibiting the elimination of synapses does lead to more neural activity. Increased ACh could be explained in a similar way – perhaps the brain neural activity that regulates the vagus nerve is also overactivated by inhibiting NAc pruning, in turn leading to more sympathetic neural activity directed to ACh-expressing T cells. Or, perhaps it’s all about the vagus! If vagal modulation of the spleen also requires noradrenergic input, then increased vagal output alone could explain an increase in both NA and ACh. There also could be a mix and match effect in which there is increased splenic nerve (NA) activity, and reduced T cell egress, resulting in increased ACh.

Why this would happen in females, but not males, is unclear.

### A link between adolescence and a broad definition of health

We have presented data indicating that development with the reward circuitry during adolescence has a sex-specific impact on the spleen. We have also shown that microglial pruning in the NAc is important for acute and persistent social outcomes [12, 39]. There are also data indicating that social history can also modulate immune cell responses to pathogens [40], suggesting that early life social context can persistently program immune responses. Social relationships and support have an inordinately impactful effect on health in a very broad sense. Strong social support promotes a variety of positive health outcomes in humans [41-43]. In rodents, social housing improves recovery after a cardiac event [44], reduces stress [45, 46], and reduces age-related memory impairments [47, 48], while social isolation increases depression- and anxiety-like behavior [49], shortens lifespan [50], and exacerbates Alzheimer’s disease pathology [51]. Thus, the data presented here would suggest that the trajectory of brain and social development during adolescence will meaningfully influence an organism’s biological response to *disease*, in a very broad sense of the word.

## Acknowledgements

We thank Dr. Justin Bourgeois for his assistance with tissue processing. This work was supported by the National Institutes of Health R01DA052889 and R03AG07011 to AMK and Albany Medical College Start-up funds to AMK.

## Author Declarations

AMK designed the experiments. JMK, IP, and AM performed the experiments. AMK analyzed the experiments. AMK wrote the manuscript. All authors edited the manuscript. The authors declare no conflicts of interest.

**Supp. Table 1:**
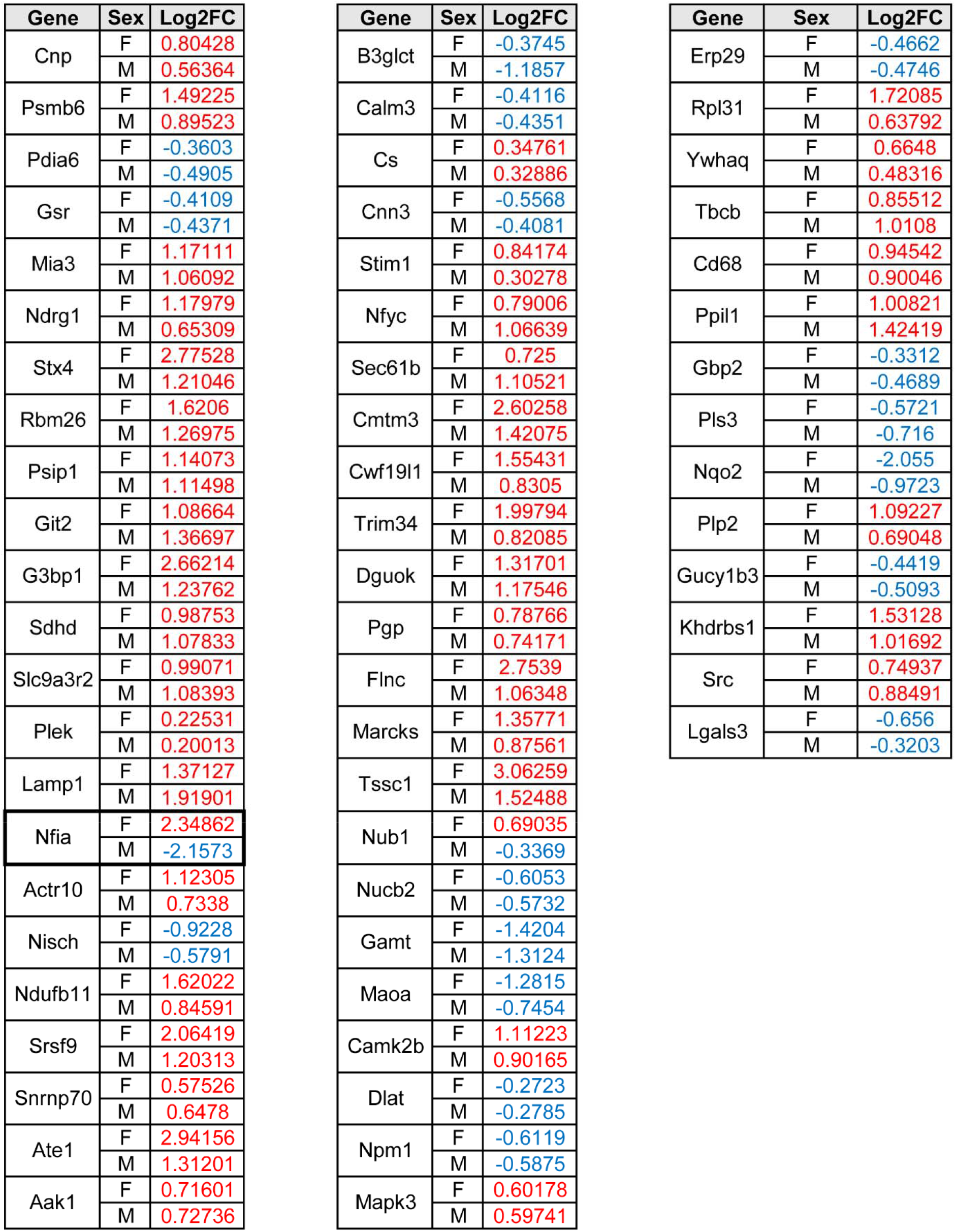
Splenic DEPs regulated by NAc pruning shared between male and female rats Only 1 of 60 splenic DEPs regulated by NAc pruning, Nfia, was not regulated in the same direction in male and female rats.

## Notes

### Competing Interest Statement

The authors have declared no competing interest.

## BIBLIOGRAPHY

1. Holt-Lunstad, J., T.B. Smith, and J.B. Layton, Social relationships and mortality risk: a meta-analytic review. PLoS Med, 2010. 7(7): p. e1000316.

2. Cacioppo, J.T. and S. Cacioppo, Social Relationships and Health: The Toxic Effects of Perceived Social Isolation. Soc Personal Psychol Compass, 2014. 8(2): p. 58–72.

3. Eisenberger, N.I., et al., In Sickness and in Health: The Co-Regulation of Inflammation and Social Behavior. Neuropsychopharmacology, 2017. 42(1): p. 242–253.

4. Kopec, A.M., C.J. Smith, and S.D. Bilbo, Neuro-Immune Mechanisms Regulating Social Behavior: Dopamine as Mediator? Trends Neurosci, 2019. 42(5): p. 337–348.

5. Snyder-Mackler, N., et al., Social status alters immune regulation and response to infection in macaques. Science, 2016. 354(6315): p. 1041–1045.

6. Spear, L.P., The adolescent brain and age-related behavioral manifestations. Neurosci Biobehav Rev, 2000. 24(4): p. 417–63.

7. Fuhrmann, D., L.J. Knoll, and S.J. Blakemore, Adolescence as a Sensitive Period of Brain Development. Trends Cogn Sci, 2015. 19(10): p. 558–566.

8. Ben-Shaanan, T.L., et al., Activation of the reward system boosts innate and adaptive immunity. Nat Med, 2016. 22(8): p. 940–4.

9. Ben-Shaanan, T.L., et al., Modulation of anti-tumor immunity by the brain’s reward system. Nat Commun, 2018. 9(1): p. 2723.

10. Schafer, D.P., et al., Microglia sculpt postnatal neural circuits in an activity and complement-dependent manner. Neuron, 2012. 74(4): p. 691–705.

11. Stevens, B., et al., The classical complement cascade mediates CNS synapse elimination. Cell, 2007. 131(6): p. 1164–78.

12. Kopec, A.M., et al., Microglial dopamine receptor elimination defines sex-specific nucleus accumbens development and social behavior in adolescent rats. Nat Commun, 2018. 9(1): p. 3769.

13. Klein, S.L. and K.L. Flanagan, Sex differences in immune responses. Nat Rev Immunol, 2016. 16(10): p. 626–38.

14. Kumar, B.V., T.J. Connors, and D.L. Farber, Human T Cell Development, Localization, and Function throughout Life. Immunity, 2018. 48(2): p. 202–213.

15. Oxford, K.L., et al., The interplay between immune maturation, age, chronic viral infection and environment. Immun Ageing, 2015. 12: p. 3.

16. Parker, G.A., et al., Histologic Features of Postnatal Development of Immune System Organs in the Sprague-Dawley Rat. Toxicol Pathol, 2015. 43(6): p. 794–815.

17. Ackerman, K.D., et al., Neonatal sympathetic denervation alters the development of in vitro spleen cell proliferation and differentiation. Brain Behav Immun, 1991. 5(3): p. 235–61.

18. Rahman, N., G. Stewart, and G. Jones, A role for the atopy-associated gene PHF11 in T-cell activation and viability. Immunol Cell Biol, 2010. 88(8): p. 817–24.

19. Clarke, E., et al., Functional characterization of the atopy-associated gene PHF11. J Allergy Clin Immunol, 2008. 121(5): p. 1148–1154 e3.

20. Cox, M.A., et al., Beyond neurotransmission: acetylcholine in immunity and inflammation. J Intern Med, 2020. 287(2): p. 120–133.

21. Nance, D.M. and V.M. Sanders, Autonomic innervation and regulation of the immune system (1987-2007). Brain Behav Immun, 2007. 21(6): p. 736–45.

22. Felten, S.Y. and J. Olschowka, Noradrenergic sympathetic innervation of the spleen: II. Tyrosine hydroxylase (TH)-positive nerve terminals form synapticlike contacts on lymphocytes in the splenic white pulp. J Neurosci Res, 1987. 18(1): p. 37–48.

23. Dantzer, R., et al., Molecular basis of sickness behavior. Ann N Y Acad Sci, 1998. 856: p. 132–8.

24. Inagaki, T.K., et al., The role of the ventral striatum in inflammatory-induced approach toward support figures. Brain Behav Immun, 2015. 44: p. 247–52.

25. McKim, D.B., et al., Sympathetic Release of Splenic Monocytes Promotes Recurring Anxiety Following Repeated Social Defeat. Biol Psychiatry, 2016. 79(10): p. 803–813.

26. Wohleb, E.S., et al., Re-establishment of anxiety in stress-sensitized mice is caused by monocyte trafficking from the spleen to the brain. Biol Psychiatry, 2014. 75(12): p. 970–81.

27. Devoino, L.V., et al., Involvement of dopamine D1 and D2 receptors in the rat nucleus accumbens in immunostimulation. Neurosci Behav Physiol, 2007. 37(2): p. 147–51.

28. Rosas-Ballina, M., et al., Splenic nerve is required for cholinergic antiinflammatory pathway control of TNF in endotoxemia. Proc Natl Acad Sci U S A, 2008. 105(31): p. 11008–13.

29. Rosas-Ballina, M., et al., Acetylcholine-synthesizing T cells relay neural signals in a vagus nerve circuit. Science, 2011. 334(6052): p. 98–101.

30. Gonzalez-Gonzalez, M.A., et al., Platinized graphene fiber electrodes uncover direct spleen-vagus communication. Commun Biol, 2021. 4(1): p. 1097.

31. Bekhbat, M., et al., Chronic adolescent stress sex-specifically alters central and peripheral neuro-immune reactivity in rats. Brain Behav Immun, 2019. 76: p. 248–257.

32. Kirkland JM, P.I., Kopec AM, Microglia-mediated synaptic pruning in the nucleus accumbens during adolescence: A preliminary study of the proteomic consequences and putative female-specific pruning target. bioRxiv, 2023.

33. Shansky, R.M., Are hormones a “female problem” for animal research? Science, 2019. 364(6443): p. 825–826.

34. Jang, N., G. Stewart, and G. Jones, Polymorphisms within the PHF11 gene at chromosome 13q14 are associated with childhood atopic dermatitis. Genes Immun, 2005. 6(3): p. 262–4.

35. Muscat, P., et al., PHF11 expression and cellular distribution is regulated by the Toll-Like Receptor 3 Ligand Polyinosinic:Polycytidylic Acid in HaCaT keratinocytes. BMC Immunol, 2015. 16: p. 69.

36. Ikari, J., et al., Plant homeodomain finger protein 11 promotes class switch recombination to IgE in murine activated B cells. Allergy, 2014. 69(2): p. 223–30.

37. Krummel, M.F., F. Bartumeus, and A. Gerard, T cell migration, search strategies and mechanisms. Nat Rev Immunol, 2016. 16(3): p. 193–201.

38. Han, D., et al., Targeting Brain-spleen Crosstalk After Stroke: New Insights Into Stroke Pathology and Treatment. Curr Neuropharmacol, 2021. 19(9): p. 1590–1605.

39. Kirkland JM, E.E., Patel I, Kopec AM, Impaired microglia-mediated synaptic pruning in the nucleus accumbens during adolescence results in persistent dysregulation of familiar, but not novel social interactions in sex-specific ways. bioRxiv, 2023.

40. Sanz, J., et al., Social history and exposure to pathogen signals modulate social status effects on gene regulation in rhesus macaques. Proc Natl Acad Sci U S A, 2020. 117(38): p. 23317–23322.

41. Bennett, D.A., et al., The effect of social networks on the relation between Alzheimer’s disease pathology and level of cognitive function in old people: a longitudinal cohort study. Lancet Neurol, 2006. 5(5): p. 406–12.

42. Atadokht, A., et al., The role of family expressed emotion and perceived social support in predicting addiction relapse. Int J High Risk Behav Addict, 2015. 4(1): p. e21250.

43. Gunnar, M.R., Social Buffering of Stress in Development: A Career Perspective. Perspect Psychol Sci, 2017. 12(3): p. 355–373.

44. Gaudier-Diaz, M.M., et al., Social influences on microglial reactivity and neuronal damage after cardiac arrest/cardiopulmonary resuscitation. Physiol Behav, 2018. 194: p. 437–449.

45. Burkett, J.P., et al., Oxytocin-dependent consolation behavior in rodents. Science, 2016. 351(6271): p. 375–8.

46. Kikusui, T., J.T. Winslow, and Y. Mori, Social buffering: relief from stress and anxiety. Philos Trans R Soc Lond B Biol Sci, 2006. 361(1476): p. 2215–28.

47. Templer, V.L., T.B. Wise, and V.R. Heimer-McGinn, Social housing protects against age-related working memory decline independently of physical enrichment in rats. Neurobiol Aging, 2019. 75: p. 117–125.

48. Morrison, K.E., et al., Peripubertal Stress With Social Support Promotes Resilience in the Face of Aging. Endocrinology, 2016. 157(5): p. 2002–14.

49. Walker, D.M., et al., Long-Term Behavioral Effects of Post-weaning Social Isolation in Males and Females. Front Behav Neurosci, 2019. 13: p. 66.

50. Razzoli, M., et al., Social stress shortens lifespan in mice. Aging Cell, 2018. 17(4): p. e12778.

51. Peterman, J.L., et al., Prolonged isolation stress accelerates the onset of Alzheimer’s disease-related pathology in 5xFAD mice despite running wheels and environmental enrichment. Behav Brain Res, 2020. 379: p. 112366.

